# High Numerical Aperture Epi-illumination Selective Plane Illumination Microscopy

**DOI:** 10.1101/273359

**Authors:** Bin Yang, Yina Wang, Siyu Feng, Veronica Pessino, Nico Stuurman, Bo Huang

**Affiliations:** Department of Pharmaceutical Chemistry, University of California in San Francisco, San Francisco, CA 94143, USA; The UC Berkeley-UCSF Graduate Program in Bioengineering, San Francisco, CA 94143, USA; Graduate Program of Biophysics, University of California, San Francisco, San Francisco, CA 94143, USA; Department of Cellular and Molecular Pharmacology, University of California, San Francisco, 600 16th Street, San Francisco, CA 94143, USA; Howard Hughes Medical Institute, San Francisco, CA 94143 USA; Department of Biochemistry and Biophysics, University of California, San Francisco, San Francisco, CA 94143, USA; Chan Zuckerberg Biohub, San Francisco, CA 94158, USA

## Abstract

**Selective-plane illumination microscopy (SPIM) provides unparalleled advantages for long-term volumetric imaging of living organisms. In order to achieve high-resolution imaging in common biological sample holders, we designed a high numerical aperture (NA) epi-illumination SPIM (eSPIM) system, which utilizes a single objective and has an identical sample interface as an inverted fluorescence microscope with no additional reflection elements. This system has an effective detection NA of > 1.06. We demonstrated multicolor and fast volumetric imaging of live cells and single-molecule super-resolution microscopy using our system.**

---

For over a decade selective-plane illumination microscopy (SPIM), or light-sheet microscopy, has been successfully used for 3D imaging applications in developmental and cell biology, anatomical science, biophysics and neuroscience^1^. Almost all light-sheet microscopes need at least two objectives close together, hence restricting the sample mounting format. In a horizontal SPIM configuration^2^ where the optical pathways are parallel to the optical table, small tubes or cylinders of agarose gel hold the sample in the space surrounded by objectives. To accommodate traditional mounting protocols such as samples prepared on glass coverslips, “dipping” configurations ^3,4^ were developed, with perpendicular optical pathways and the objectives pointing downwards. On the other hand, an “open-top” configuration with the objectives pointing upwards ^5^–^7^ potentially allows a SPIM system to be operated like an inverted fluorescence microscope and accept conventional biological sample formats including multi-well plates. Among such methods, Oblique Illumination Microscopy^8^ and Swept Confocally Aligned Planar-Excitation microscopy (SCAPE)^9^ use a single objective lens for illumination and detection without additional reflecting elements^10^–^12^ in the sample space. The sample is illuminated obliquely, resulting in a tilted illumination plane. This tilting is corrected by a remote imaging module in the detection path so that all light arrives on the camera in focus. However, the remote imaging module leads to a loss of numerical aperture (NA), such that both systems have a NA < 0.7^8,9^, whereas a high NA is essential to achieving the resolution for subcellular imaging and sensitivity for single-molecule detection.

Here, we designed a single-objective oblique epi-illumination SPIM (eSPIM) system to solve the problem of limited detection NA (Fig. 1A, Supplementary Figs. S1 and S2). In this design, a water-immersion objective (O1) of NA 1.27 is used for both illumination and fluorescence collection. The excitation light sheet has an incident angle of 60° relative to the optical axis of O1, with an effective excitation NA of ∼0.3 and a waist and length of ∼1 µm and ∼12.8 µm,respectively. The remote imaging module contains two objective lenses (O2 and O3) arranged at an angle of 30°, so that the intermediate image produced by O2 is re-imaged by O3 in focus. This angled arrangement was exactly the cause of the NA loss previously, because it shifts part of the light cone generated by O2 outside of the collectable range of O3. When the NA of O2 is high enough to ensure sufficient coverage of the NA of O1, it is impractical for O3 to have an even larger collection cone angle. To solve this problem, we chose a mismatched pair of objectives for the remote imaging module: an air objective for O2 (NA = 0.9) and a water-immersion objective for O3 (NA = 1.0) (Supplementary Fig. S3). A 3D-printed water container (Fig. 1A) separates the focal space of the two objectives by a piece of coverglass at the intermediate image plane, with one side being air, the other side being water, and the z’ positon of the coverglass adjusted to minimize the spherical aberration. The refractive index difference between the working media of O2 and O3 compresses the angle of the O2 light cone, thus minimizing NA loss. By mounting all components of the remote imaging module on the same translation stage, we found that their alignment is robust and stable, and routine realignment is unnecessary.

**Figure 1.**
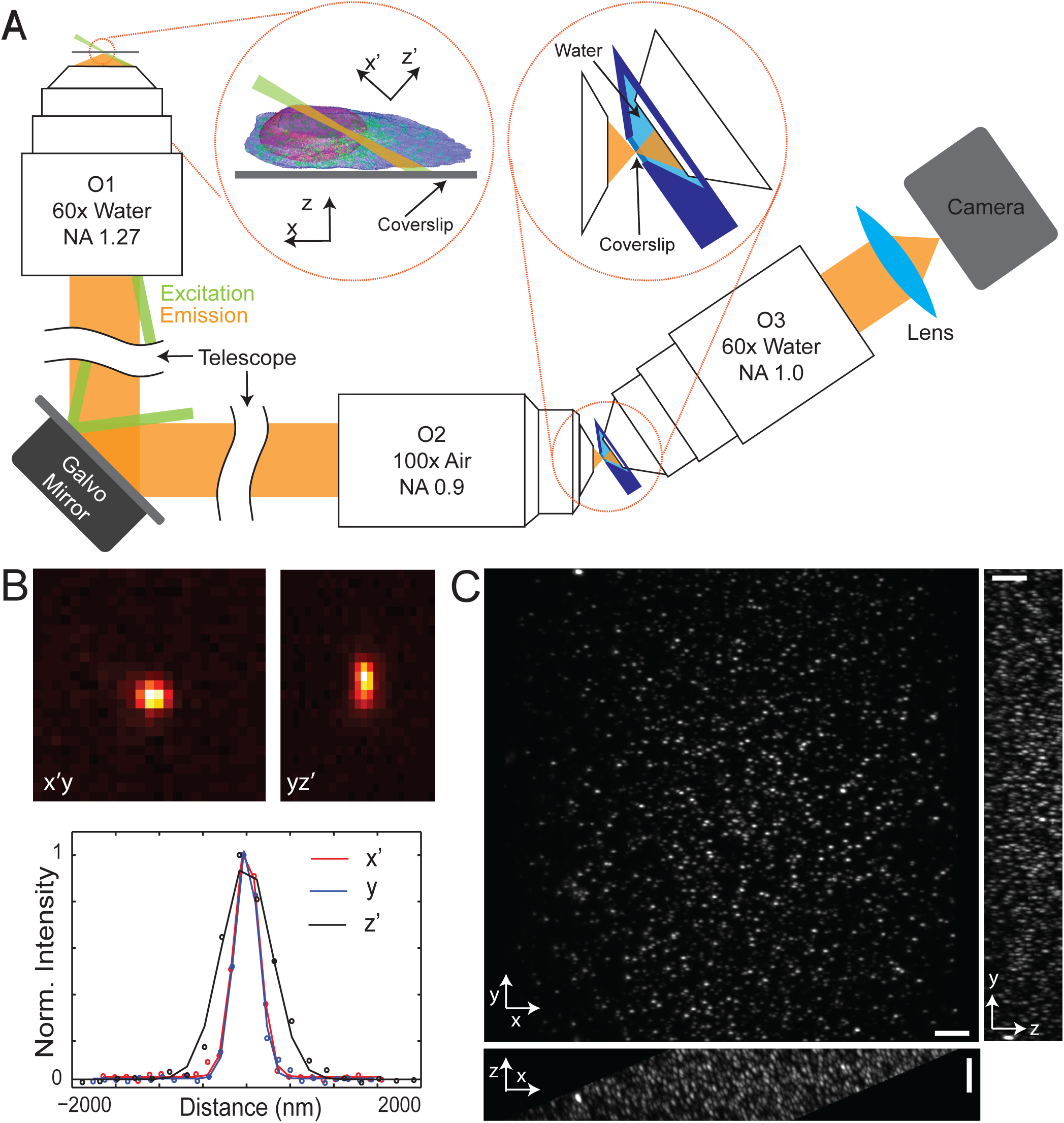
Optical setup and characterization of resolution and field-of-view. (**A**) Scheme of the experimental setup. O1 is used for both illumination and fluorescence detection. At the focal space of O1, the excitation light (shown in green) generates an oblique light sheet, 60° relative to the optical axis. A remote imaging module composed by two objective lens O2 and O3 is essential to imaging the oblique plane in focus. A water container in the focal space of O2 and O3 separates them by a glass coverslip, with one side being air medium and the other side water. Left inset shows the definition of the coordinate system. Right inset shows the detailed design of the remote imaging module. (**B**) PSF of the eSPIM, measured with 45 nm fluorescent green beads. Representative cross-sections of the PSF in the x’y-plane, in the yz‘-plane, intensity plots along the three axes of a bead are shown. (**C)** Maximal intensity projection of the bead image in different planes. The averaged FWHMs are 316 nm, 277 nm and 850 nm along the x‘-, y-and z‘-axes, respectively, calculated for all beads within the field of view. Scale bar is 5 µm.

Imaging an oblique plane requires obtaining a perfect (i.e. aberration-free) intermediate image of a volume. Optical systems simultaneously satisfying both the sine and Herschel conditions fulfill this requirement^13^. To achieve these conditions, the pupil planes of O1 and O2 are conjugated, and the lateral magnification from the sample space to the intermediate image is set to be 1.33, which is the ratio between the refraction indexes of the working media of O1 and O2. Under this condition, the axial magnification is also 1.33. The effective detection NA of our system is estimated to be ∼ 1.20 along the y axis and ∼ 1.06 along the x‘-axis (Supplementary Fig. S3). Our optical simulation further confirmed that the performance of our system is consistent within a volume of 70 µm × 70 µm × 20 µm (Supplementary Fig. S4). The overall transmission efficiency of the remote imaging module is 73% (see Methods). To obtain a volumetric image, a Galvo mirror conjugated to the pupil planes of both O1 and O2 scans the excitation light sheet and descans the image so that the intermediate image is always projected at the focal plane of O3. Neither the microscope stage nor the objective O1 need to move, ensuring mechanical stability. The entire system is controlled by the open-source software of Micro-Manager.

To measure the resolution of our system, we imaged 45 nm fluorescent blue beads (Figs. 1B and 1C). The full-widths at half-maximums (FWHMs) of the bead images were 316 nm, 277 nm and 850 nm along the x‘-, y-and z‘-axes, respectively, matching our detection NA estimation. Within an imaging volume of about 70 µm along the y direction and 20 µm along the z direction (Fig. 1C), the aberration was negligible. This imaging depth is similar to that reported by Lattice Light Sheet Microcopy^4^. The Galvo mirror faithfully scans the oblique light sheet in the x direction for ∼ 100 µm, which can be further increased by scanning the sample stage if necessary.

To demonstrate the performance of our system in live cell microscopy, we imaged a variety of subcellular structures in cell lines grown in 8-well coverglass-bottom chambers, including microtubules and mitochondria in HeLa cells (transient over expression of EGFP-tubulin-6 and mRuby2-TOMM20-N-10, respectively) (Supplementary Figs. S5 and Movies 1-2), endogenously labeled clathrin structures in HEK293T cells (mNeonGreen2_11_ knock-in for CLTA) (Fig. 2A and Supplementary Movie 3-4), as well as two-color imaging of nuclear lamina and lysosomes in HEK293T cells (mNeonGreen2_11_ knock in for LMNA and LysoTracker Deep Red staining, respectively) (Fig. 2B and Supplementary Movies 5-6). The puncta of clathrin structures showed similar dimensions as measured earlier with fluorescent beads, demonstrating the practical spatial resolution of our system and enabling tracking of the movement of clathrin structures across the entire volume of cells. Even for endogenously labeled proteins, which typically exhibit much lower fluorescence signal than over-expressed ones, we were able to acquire live cell movies at 0.5-2 volumes per second continuously for more than 10 minutes. We verified the greatly-reduced photobleaching of our system compared to a spinning disk confocal microscope when recording a similar level of fluorescence signal from the same sample (Supplementary Fig. S6).

**Figure 2.**
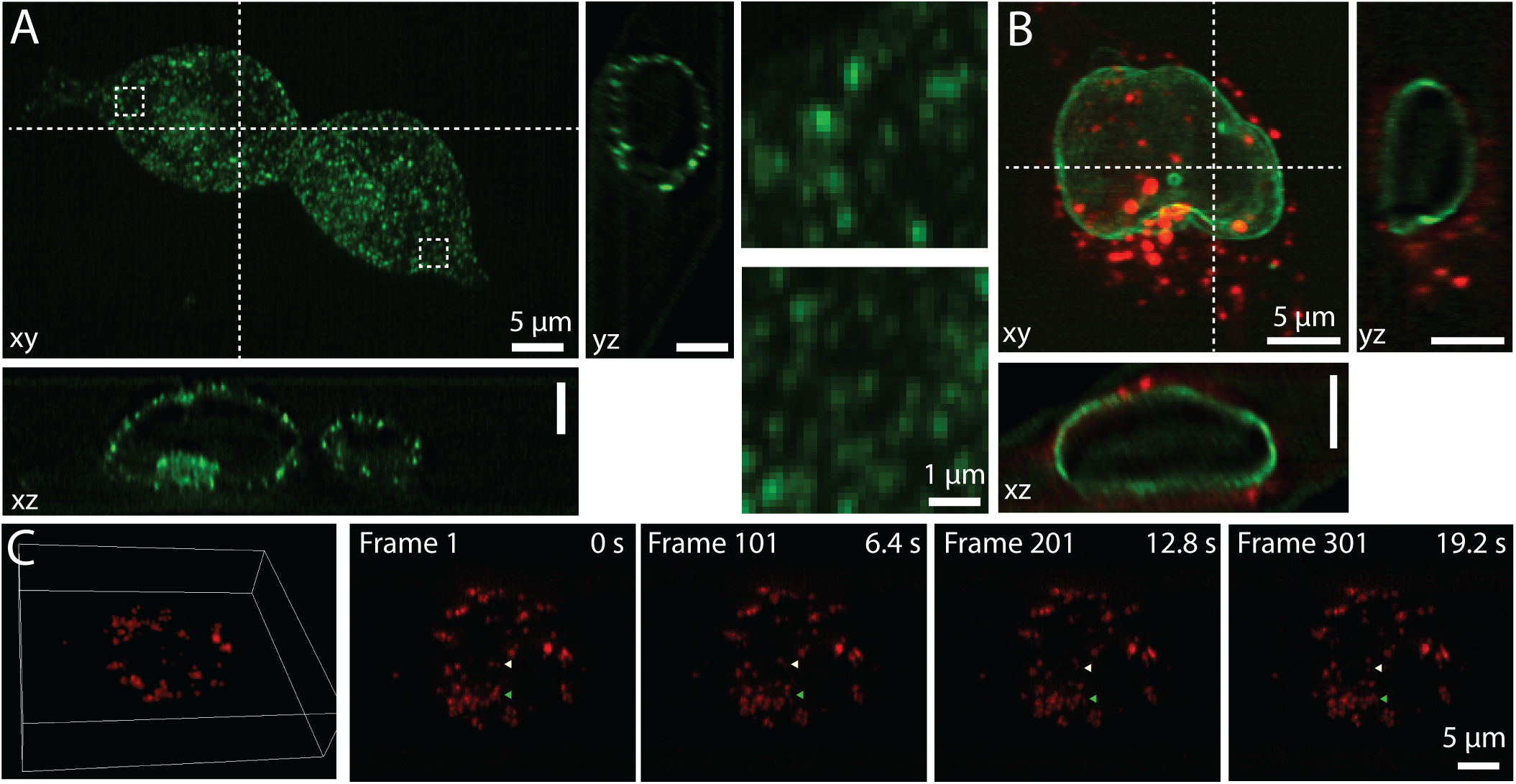
Volumetric live imaging of mammalian and *Drosophila* S2 cells. (**A**) Two HEK293T cells with endogenous clathrin A labeled by mNeonGreen2_11_ knock-in. (**B**) A HEK293T cell with endogenous lamin A/C labeled by mNeonGreen2_11_ knock-in (green) and lysosomes stained by LysoTracker Deep Red dye (red). The maximum intensity projection in the x-y plane and representative x-z and y-z slices along the dashed lines are shown. In **A**, the regions out lined bythe dashed rectangles are also shown as magnified images. (**C**) A *Drosophila* S2 cell with lysosomes labeled with LysoTracker Deep Red dye. From left to right: one 3D view image and four maximum intensity projection images at different time points. The triangles highlight two noticeable events. Images were acquired at 15 volumes per second. See Supplementary Movies 3-9.

Our system is particularly suitable for fast volumetric imaging because the Galvo mirror is the only moving mechanical element in the system; thus, the scanning speed can be pushed to the limit of the camera’s readout. To demonstrate this capacity of fast imaging, we imaged *Drosophila* S2 cells with lysosomes labeled with LysoTracker Deep Red at 14.7 volumes per second (volume size 35 µm × 35 µm × 7 µm, 34 slices per volume and camera running at 500 frames per second). At this imaging speed, we could reliably perform 3D tracking of lysosome movement dynamics (Fig. 2C and Supplementary Movies 7-9). This imaging speed is ∼ 4-5 times faster than other high-resolution light-sheet microscopes reported^4,14^. This fast imaging capability will be extremely useful for live cell single-particle tracking, calcium imaging, etc.

The high detection NA of our system enables single-molecule imaging. As a demonstration, we performed single-molecule-switching-based super-resolution microscopy for HEK293T cells stained for Lamin A/C (Supplementary Fig. S7). On average > 2000 photons were detected for each photoswitching event when imaged at 50 frames per second. The oblique light sheet restricts excitation to a selective plane, reducing out-of-focus bleaching and out-of-focus background. Therefore, our system can be particularly useful for the imaging of densely labeled cells or with point-accumulation-for-imaging-in-nanoscale-topography (PAINT)^15^.

Our microscope design provides a versatile platform to image live samples at high spatial-temporal resolution. As a unique advantage, it can be built as an add-on unit to existing inverted fluorescent microscopes (in a similar manner as a confocal spinning disk unit), converting a high-NA epifluorescence microscope into a SPIM system without modifying the sample stage. It can be easily integrated with wide-field epi-fluorescence microscopy for optogenetics and FRAP experiments. Our design is also inherently compatible with numerous methods to improve the performance of light-sheet microscopy, including various excitation schemes such as digitally scanned light sheet (to obtain more uniform illumination), modulated excitation such as Bessel^16,17^ and Airy^18^ beams (to achieve thinner light sheet across a larger view field), the use of adaptive optics^6,19^ to reduce the systematic aberration, and the application of multi-view imaging^20^ to obtain isotropic resolution.

## Author contributions

B.Y. designed and built the microscope and performed the simulations and the experiments. B.Y., Y.W., S.F. and V.P. prepared the cell samples. B.Y. and N.S. implemented the Micro-Manager^21^ software for device control and wrote custom script for fast data acquisition. B.Y.,Y.W. and B.H. analyzed the data. B.H supervised the project. B.Y and B.H wrote the manuscript.

## Acknowledgements

We acknowledge Tom Goddard and Thomas Ferrin from UCSF for their help in using ChimeraX. We thank Edaeni Hamid from Nikon for her help in providing the technical information of the objective lenses. This project is supported by National Institutes of Health (R33EB019784). B.H. is a Chan Zuckerberg Biohub investigator.

## Conflict of interests

A provisional patent application has been filed covering the reported microscope design.

## Methods

### Optical setup

A water-immersion objective (O1, Nikon CFI Plan Apo IR 60XWI) of NA 1.27 was used for both illumination and fluorescence collection. The illumination light came from three lasers (Votran Stradus 488nm and 642nm, Coherent Sapphire 561nm). The beams were combined by two dichroic mirrors, collimated by a telescope composed of two achromatic lens, expanded by two cylindrical lens (Thorlabs CL 50mm and CL 200mm) to form an elongated spot, and then clipped by a mechanical slit conjugated with the pupil plane of O1. The illumination beam was then reflected by a dichroic mirror (DM, Chroma ZT405/488/561/640rpc) and intersected the back aperture of O1 off-center to generate an oblique light sheet at the focal space of O1. The remote imaging module consisted of two objective lenses (O2, Nikon CFI LU Plan Fluor EPI P 100x and O3, Nikon CFI Fluor 60XW). The pupil planes of O1 and O2 were conjugated by two 4f-systems (L1-L4). The optical axis of O3 was 30° relative to that of O2 so that O3 could re-image the intermediate image in focus. A 3D-printed water container separated the focal space of the two objectives by a glass coverslip, with one side being air and the other side water. The water container was mounted on a motorized translation stage (Thorlabs PT1-Z8) so that it could be translated along the optical axis of O3. The objective lens O1 was mounted on a manual translation stage (Thorlabs CT1) for focus adjustment. The objective lens O3 was mounted on a piezo stage (Thorlabs DRV517) so that its focus could be finely tuned. The fluorescence was filtered by either individual band-pass filters (Chroma ET525/50m, ET605/70m and ET705/72m) or a quad-band filter (Chroma ZET405/488/561/640x) and then detected by a scientific CMOS camera (PCO Edge). The pixel size of the camera at the sample space was 133 nm.

A galvo mirror (Thorlabs, GVS011) was conjugated to both the pupil planes of O1 and O2. Rotating the Galvo mirror scans the oblique light sheet across the sample (along the x-axis), with the incident angle kept as 60°. The galvo mirror also descans the intermediate image at the focal space of O2. The scanning frequency of the galvo mirror can be as fast as a few hundred Hz. Hence, the imaging speed was mostly limited by the readout time of the camera and the power of the excitation laser. The frame rate of the camera is as fast as 800 frames per second for a region of interest of 640 × 256 pixels. The galvo mirror scans the light sheet faithfully across ∼ 70 µm. Out of this range, either the illumination or the fluorescence light starts to be cropped. It is also possible to scan the sample with the microscope stage (PI, PILine M-687.UN) for longer ranges, albeit at slower speed.

To measure the transmission efficiency of the remote imaging module, collimated laser light was let to pass through the component. The transmission efficiency was then obtained by calculating the ratio of the exiting and entering light power, being respectively 72.7% at 488 nm, 66.7% at 561 nm and 62.3% at 641 nm.

### Characterization of resolution with fluorescent beads

45 nm fluorescent beads were imaged to characterize the resolution of the system. The beads were firstly embedded in 2% agarose gel and then sandwiched between a glass coverslip and a glass slide. The sample was then placed on the microscope stage with the coverslip side facing the objective. The wavelength of excitation light was 488 nm and the corresponding emission filter is a bandpass 525/50 filter. The raw images were slices across the sample in the x‘-y plane, 30° to the x-y plane. Figure 1B shows the cross-sections of a bead image in the x’y-and yz‘-planes. The FWHMs of the intensity plots gave the resolution to be respectively 316 nm, 277 nm and 850 nm along the x‘-, y-and z‘-axes. These values were obtained by averaging the FWHMs of all beads in the field of view.

### Data acquisition, processing and viewing

Micro-Manager^21^ was used for device control and multi-dimensional data acquisition. The galvo scanner was set up as a DA (digital-analogue)-z stage, controlled by one Analog Output channel of the NI PCIe 6323 DAQ card. The lasers emission states were controlled via an Arduino Uno board. The galvo scanner and the lasers were hardware-synchronized^21^ through the TTL output of the PCO sCMOS camera.

The raw SPIM data were obtained in the x‘-y-z’ space (see Figure 1A). The data were then descrewed, deconvolved by the measured PSF and rotated to the x-y-z space. This process was performed with a free-online package (https://www.flintbox.com/public/project/31374/) produced by Janelia Research Campus. Deconvolution was applied to the cell images shown in Figure 2 (same as the previously used image processing procedure for Lattice Light Sheet microscopy^4^), but not to the bead image for resolution measurement in Figure 1

The free software ChimeraX^22^ by UCSF was used to view and demonstrate the volumetric date in 4D.

### Global Exposures with Rolling Shutter

The rolling shutter mode of our sCMOS camera provides readout time as short as 1 ms for a region of interest of 200 rows, facilitating fast imaging. Although this mode is fast, the readout of each row is no longer simultaneous. When the frame rate is close to the maximum readout speed of the camera, this asynchronous readout causes PSF distortion (Supplementary Fig. S8). Although global shutter mode can solve this problem, it increases the readout noise and slows down the readout by a factor of 2. Therefore, we implemented global exposure with rolling shutter by triggering pulsed laser illumination only during the time when all rows were exposing (Supplementary Fig. S8). The effective imaging time for each frame was thus the exposure time plus the readout time. Global exposure was realized through the NI PCIe 6323 DAQ card using LabVIEW programming.

### Cell culture and transfection

The conditions for cell culture, transfection of HEK 293 T and HeLa cells were described in our previous publication^23^. The knock-in cell lines, including the sequences for the guide RNAs and donor DNAs, were previously generated in the same publication^23^. To prepare sample for microscopy, knock-in cells were grown on an 8-well glass bottom chamber (Thermo Fisher Scientific). In order to achieve better cell attachment, 8-well chamber was coated with fibronectin (Sigma-Aldrich) for one hour before seeding cells.

*Drosophila* S2 cells were cultured in Schneider’s Drosophila Medium (Gibco) supplemented with 10% heat inactivated fetal bovine serum and penicillin/streptomycin (50 µg/ml). The cells were plated into 8-well plates coated with 0.5 mg/mL solution of Concanavalin A and then incubated for 1 day. The next day 225 µL 50 nM LysoTracker Deep Red dye solution was added to each well for 30 minutes prior to imaging.

SpyTag-LaminA/C cells were prepared as previously described^24^. Briefly, HEK293T cells were plated in 8-well chambers, and left to adhere for 24 hours. Before plating, chambers were coated with Poly-L-Lysine (Sigma) for 20 minutes, and washed three times with phosphate buffered saline (PBS). At 24 hours, cells were transfected with 100ng per well of SpyTag-LaminA/C. At 48 hours post-plating, cells were fixed with 2% Paraformaldehyde for 30 minutes at room temperature, followed by three washes with PBS. Samples were then blocked and permeabilized with 3% Bovine Serum Albumin (BSA) and 2% NP40 for one hour at room temperature, followed by over-night room-temperature incubation with SpyCatcher-Alexa647 (80 nM in 3% BSA). Finally, samples were washed 5 times with PBS.

